# Stimulation Effects Mapping for Optimizing Coil Placement for Transcranial Magnetic Stimulation

**DOI:** 10.1101/2024.02.23.581706

**Authors:** Gangliang Zhong, Fang Jin, Liang Ma, Yongfeng Yang, Baogui Zhang, Dan Cao, Jin Li, Nianming Zuo, Lingzhong Fan, Zhengyi Yang, Tianzi Jiang

**Affiliations:** Brainnetome Center, Institute of Automation, Chinese Academy of Sciences, Beijing 100190, China; Shanghai Mental Health Center, Shanghai Jiao Tong University School of Medicine, Shanghai 200030, China; Department of Psychiatry, Henan Mental Hospital, The Second Affiliated Hospital of Xinxiang Medical University, Xinxiang 453002, China; Henan Key Lab of Biological Psychiatry, Xinxiang Medical University, Xinxiang 453002, China; International Joint Research Laboratory for Psychiatry and Neuroscience of Henan, Xinxiang 453002 China; State Key Laboratory of Brain and Cognitive Sciences, Beijing MRI Center for Brain Research, Institute of Biophysics, Chinese Academy of Sciences, Beijing, China; University of Chinese Academy of Sciences, Beijing, China; National Laboratory of Pattern Recognition, Institute of Automation, Chinese Academy of Sciences, Beijing 100190, China; Center for Excellence in Brain Science and Intelligence Technology, Institute of Automation, Chinese Academy of Sciences, Beijing 100190, China; School of Artificial Intelligence, University of Chinese Academy of Sciences, Beijing 100190, China

**Keywords:** Transcranial magnetic stimulation, E-field, stimulation effects mapping, coil placement

## Abstract

**Background:** The position and orientation of transcranial magnetic stimulation (TMS) coil, which we collectively refer to as coil placement, significantly affect both the assessment and modulation of cortical excitability. TMS electric field (E-field) simulation can be used to identify optimal coil placement. However, the present E-field simulation required a laborious segmentation and meshing procedure to determine optimal coil placement.

**Objective:** We intended to create a framework that would enable us to offer optimal coil placement without requiring the segmentation and meshing procedure.

**Methods:** We constructed the stimulation effects map (SEM) framework using the CASIA dataset for optimal coil placement. We used leave-one-subject-out cross-validation to evaluate the consistency of the optimal coil placement and the target regions determined by SEM for the 74 target ROIs in MRI data from the CASIA, HCP15 and HCP100 datasets. Additionally, we contrasted the E-norms determined by optimal coil placements using SEM and auxiliary dipole method (ADM) based on the DP and CASIA II datasets.

**Results:** We provided optimal coil placement in ‘head-anatomy-based’ (HAC) polar coordinates and MNI coordinates for the target region. The results also demonstrated the consistency of the SEM framework for the 74 target ROIs. The normal E-field determined by SEM was more significant than the value received by ADM.

**Conclusion:** We created the SEM framework using the CASIA database to determine optimal coil placement without segmentation or meshing. We provided optimal coil placement in HAC and MNI coordinates for the target region. The validation of several target ROIs from various datasets demonstrated the consistency of the SEM approach. By streamlining the process of finding optimal coil placement, we intended to make TMS assessment and therapy more convenient.

## Introduction

Transcranial magnetic stimulation (TMS) is an effective and non-invasive technique to assess and modulate cortical excitability (McClintock et al., 2018; Ni and Chen, 2015; Walsh and Cowey, 2000). The assessment of cortical excitability can be conducted by single-pulse TMS (spTMS). Repeated TMS (rTMS) was believed to regulate cortical excitability and has promising therapeutic uses in the treatment of mental conditions such as depression (Philip et al., 2019). Assessment and treatment outcomes were both significantly impacted by the position and orientation of TMS coil, which we collectively called coil placement (Dannhauer et al., 2024). Therefore, determining optimal coil placement was crucial in research and therapeutic settings, whether spTMS or rTMS was used.

The electric field (E-field) simulation, which demonstrated TMS effects on the cortex, has been employed to optimize the placement of TMS coil. Gomez-Tames et al. (2018) showed that utilizing effective E-fields could be advantageous in identifying the optimal coil placement. In order to focus the research on a target region, the E-field was computed inside the region of interest (ROI) in the cortex. Gomez et al. (2021) have noted that a rapid computational auxiliary dipole method (ADM) could accelerate the process of optimizing coil placement. Their research significantly decreased the duration of E-field simulations of coil placements (15 minutes for 1 million simulations) for the Ernie head model in SimNIBS 3.1. However, when testing for the new subjects, it was inevitable to perform the process of segmentation and meshing for the new subjects’ MRIs when determining optimal coil placements. The segmentation and meshing offered by SimNIBS, such as headreco or mri2mesh, required a duration beyond two hours (Nielsen et al., 2018; Windhoff et al., 2013). If new technologies could be used to obtain prior knowledge for optimal coil placement, it would be possible to streamline the process by eliminating segmentation and meshing.

The brain atlas was created as prior knowledge to map the structural and functional properties of the brain by using a template dataset (Fan et al., 2016; Toga and Mazziotta, 2002). We were inspired by the brain atlas to create an innovative framework that used template dataset to give the prior knowledge for optimal coil placement. In the framework, not only the magnitude but the focality of the E-field were considered as optimization factors for the coil placement. Furthermore, Xiao et al. (2018) developed the transcranial brain atlas, based on which we proposed an updated head anatomy coordinate system with improved cross-subject reproducibility. This advancement enabled us to locate the optimal coil placement on the scalp.

In this paper, we proposed the Stimulation Effects Mapping (SEM) framework (Figure 1) to identify optimal coil placement. In the SEM framework, first, we calculated the E-field for the large-scale coil placements, which transformed to a polar coordinate system. Then, the intensity-focality value (IFV) of E-field was used as the factor to define optimal coil placements. We validated the SEM consistency on independent datasets. Finally, we contrasted the E-norms calculated by the optimal coil placements using SEM with those obtained through ADM. Our aim was to bypass the segmentation and meshing stages to provide optimal prior knowledge of TMS coil placement. This would make TMS assessment and treatment easier and more accurate.

**Figure 1.**
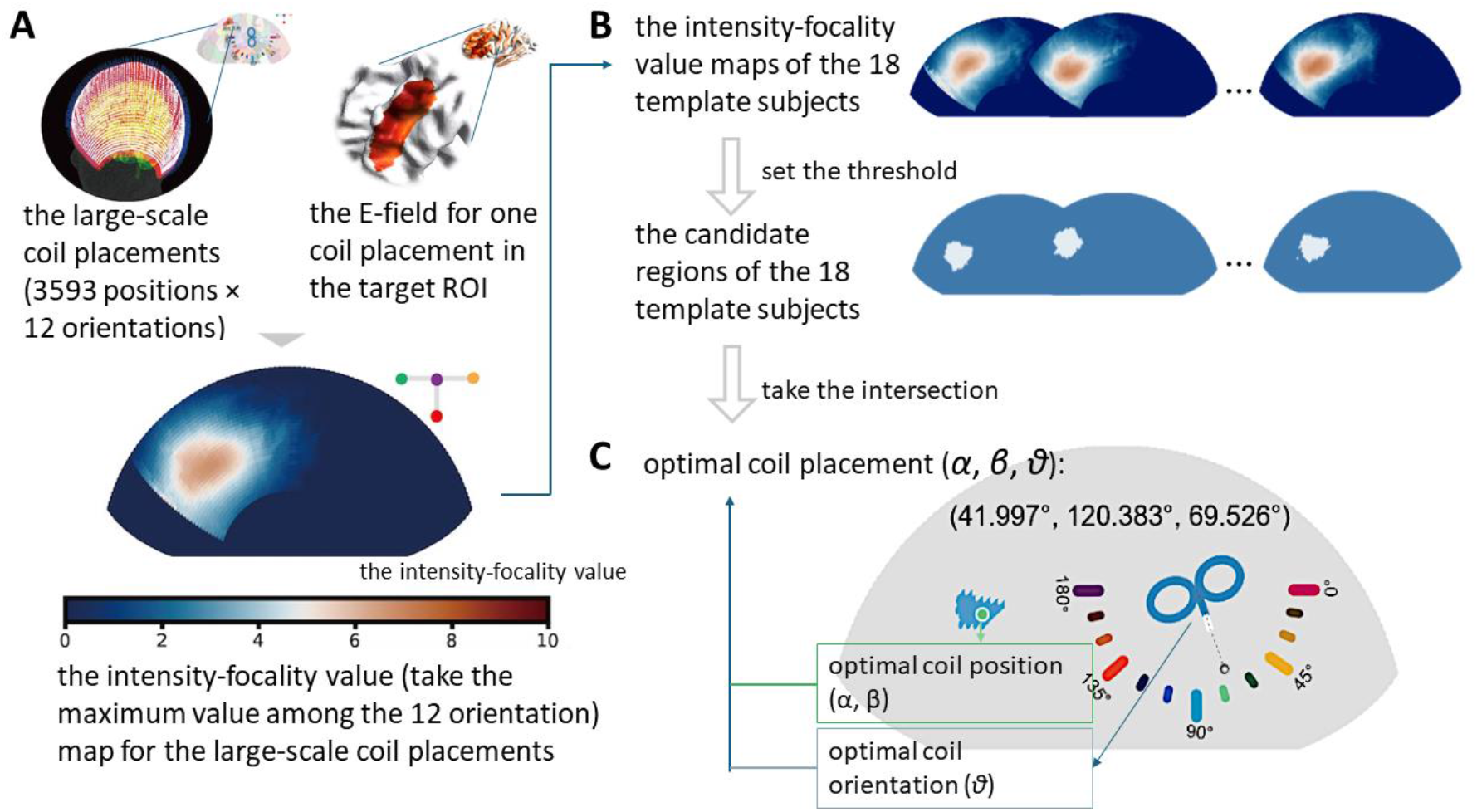
Flow chart of the SEM framework. Panel A outlines the process of calculating the intensity-focality value (IFV) map of the E-field for large-scale coil placements. Panel B describes the step of selecting potential regions from a group of 18 template subjects. Panel C explains the method for optimal coil placement.

## Materials and Methods

### Datasets

The MRI data of 18 healthy subjects (9 males, aged 25.28 ± 2.05 years) were obtained from the CASIA dataset and served as the template dataset to build the SEM framework. To evaluate the SEM framework, we analyzed the MRI data obtained from the four other datasets: HCP15, HCP100, DP and CASIA II. Detailed explanations of the procedure for obtaining MRI data for each dataset can be found in Datasets & Table S1 & Figure S9 of *Supplementary Material*. The Ethical Committee of the Institute of Automation, Chinese Academy of Sciences approved the study.

### SEM framework

Figure 1 illustrates the SEM framework. First, in Figure 1A, the large-scale coil placements (including 3593 positions × 12 orientations) for all 18 template subjects in the CASIA dataset were standardized based on a ‘head-anatomy-based’ polar coordinate (HAC) system (see Figure S11 in *Supplementary Material* for more details). These coil positions were defined with a 1.8-degree interval of *α*and *β*, and an approximate 2 mm Euclidean distance between adjacent positions. Additionally, we considered 12 different coil orientations with a 15-degree interval of *θ*, covering a total of 180 degrees on the scalp (Figure S12). We simulated the E-field of every coil placement in the cortex using the SimNIBS software (Thielscher et al., 2015) (*Supplementary Method S1*).

To confine the TMS E-field to a concentrated region, we defined the target ROI as the overlapping region between the boundary of A9/46v (ventral area 9/46) region of the middle frontal gyrus (Fan et al., 2016) and a sphere with a diameter of 10 mm. The center of the sphere and the center of the boundary were coincident. The results for each step were shown in Figure S1 to S4 of the *Supplementary material*. The computational cost for the E-field simulations for 43,116 coil placements was approximately 4,000 cores, with each subject taking 4.5 hours. We utilized 2.5 GHz 2000 Hygon C86 cores, each equipped with 6 GB of RAM.

The intensity-focality value (IFV) map for a specific template subject with the large-scale coil placements was shown in Figure 1A’s bottom image. The color gradient ranging from blue to red represented different degrees of maximum IFV for the specific orientation of the coil positions. The IFV for E-filed of every coil placement *E(α, β, θ)* was determined by Formula 1.

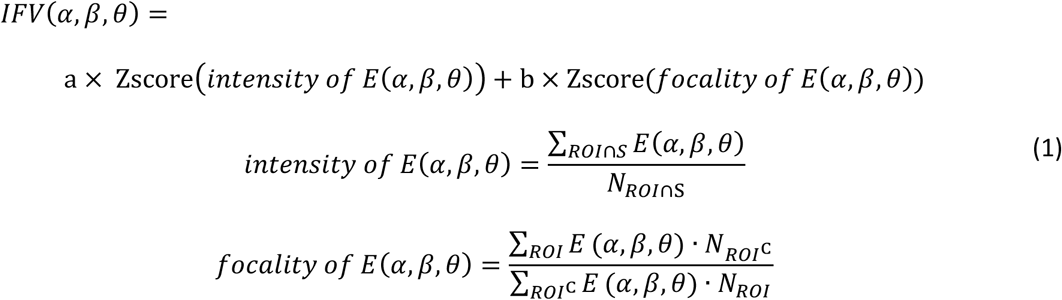

The parameters a and b were utilized to modify the weight of intensity or focality. In this article, the values of a and b were both 1. The *ROI* corresponding to the target region, while the *Sphere* was a set of tetrahedrons located within the range of the 10 mm sphere. *N*_*ROI*_ was the number of tetrahedrons in the *Sphere* and *E(α, β, θ)* was the absolute value of the E-field at each tetrahedron for each coil placement. If a collection of tetrahedrons within the range of grey matter were defined as a universal set 𝕌, then *ROI*^c^ was the complement of the ROI set. To maintain consistency in the scale of intensity and focality of E-field, the *intensity* and *focality of E(α, β, θ)* was normalized using a Z-score.

Figure 1B and C illustrate the procedure for optimal coil placement. We identified the candidate regions where the IFV exceeded 0.85 maximum IFVs for the 18 template subjects to limit the selected region. Then, we identified the region *s* that was intersected by the candidate regions of the 18 template subjects. Finally, the optimal coil placement *(α*_*optimal*_, *β*_*optimal*_, *θ*_*optimal*_) was defined in Formula 2.

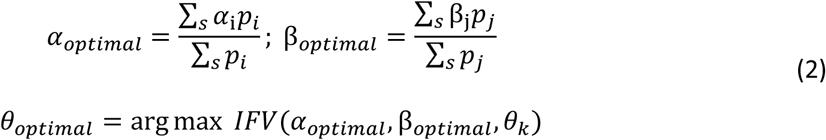

In the formula, *p* represented the population probability for every coil placement in the region *s*, which consisted of *i* coordinate *α*_*i*_, *j* coordinate *β*_*j*_, and *k* coil rotations *θ*_*k*_. In addition to finding the optimal coil placement for one specific ROI, we also tried to identify the coil placements at the group level for more than 74 target ROIs (see *Supplementary Method S2* for more details).

### Statistics

Leave-one-subject-out cross-validation was used in CASIA, HCP15, and HCP100 datasets to test the consistency of the results from the SEM framework for the 74 target ROIs. In leave-one-subject-out cross-validation, the results were determined by analyzing data from n-1 template subjects for each dataset, including data from n subjects. We calculated the Dice coefficients for the candidate regions and Euclidean distances for optimal coil placements in all 74 target ROIs. Leave-one-subject-out cross-validation was also used to validate the coil position and orientation differences in the datasets of CASIA, HCP15 and HCP100.

The E-field normal (E-norm) values were calculated based on the optimal coil placements determined from the SEM and ADM approaches using DP and CASIA II datasets. We employed the Kruskal-Wallis H-test to analyze the Z-scored average E-norm value differences among the different approaches. Dunn-Bonferroni correction was necessary. Statistics were performed using several Python packages, including scipy, sklearn, and statsmodels. For these non-parametric analyses, we employed Python packages such as scipy.stats.mannwhitneyu and statsmodels.stats.multitest. The significance threshold was determined at α = 0.05.

## Results

The optimal coil placement for the target region was determined to be at HAC coordinates (41.997°, 120.383°, 69.526°) (Figure 2B), which corresponded to MNI coordinates of −60 mm, 56 mm, and 27 mm. The mean and standard deviation of IFVs and optimal coil orientations around the optimal coil position were shown in Figure 2C. The average IFV was 5.470 ± 0.132, and the average optimal orientation was 110.554° ± 8.938°.

**Figure 2.**
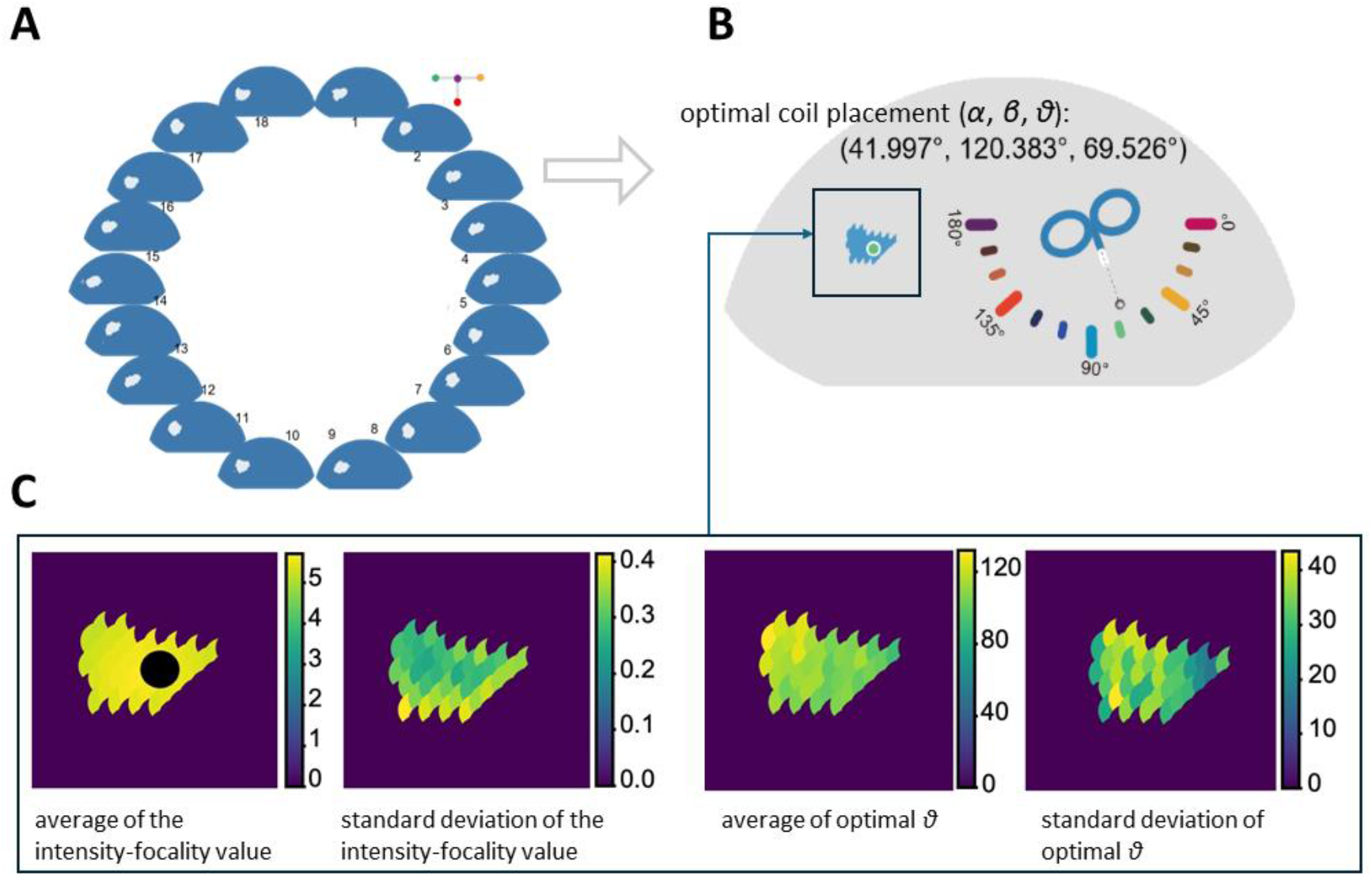
Optimal coil placements identified by SEM. A: the candidate regions for the target region in all subjects (*n* = 18) in the CASIA dataset. B: the optimal coil placement for the target region. C: the mean and standard deviation of the IFV and the optimal coil orientation, respectively. The black circle in the first figure represents optimal coil placement.

We have also expanded the SEM calculation from one target ROI to several ROIs and across many datasets for validation. Figure 3A displays the candidate regions of the 74 target ROIs in the CASIA dataset. The top image in Figure 3B depicts the Dice coefficients of the candidate regions for the 74 target ROIs. This value was cross-validated using the leave-one-subject-out approach on the CASIA dataset. The bottom image in Figure 3B represents the Euclidean distances between the optimal coil position and the nearest site on the scalp for the 74 target ROIs based on the leave-one-subject-out approach. In addition, we employed CASIA, HCP15 and HCP100 datasets to validate the SEM. In Figure 3C, we computed the mean values and standard deviations of the coil position and orientation differences between the 74 target ROIs using the leave-one-subject-out method. We calculated the Dice coefficients for the candidate regions in the 74 target ROIs across the three datasets. The Dice coefficients, computed for all 74 ROIs, were 0.806 ± 0.144 for comparing the CASIA and HCP100 datasets. They were 0.519 ± 0.247 for the comparison between the CASIA and HCP15 datasets and 0.599 ± 0.216 for the comparison between the HCP15 and HCP100 datasets.

**Figure 3.**
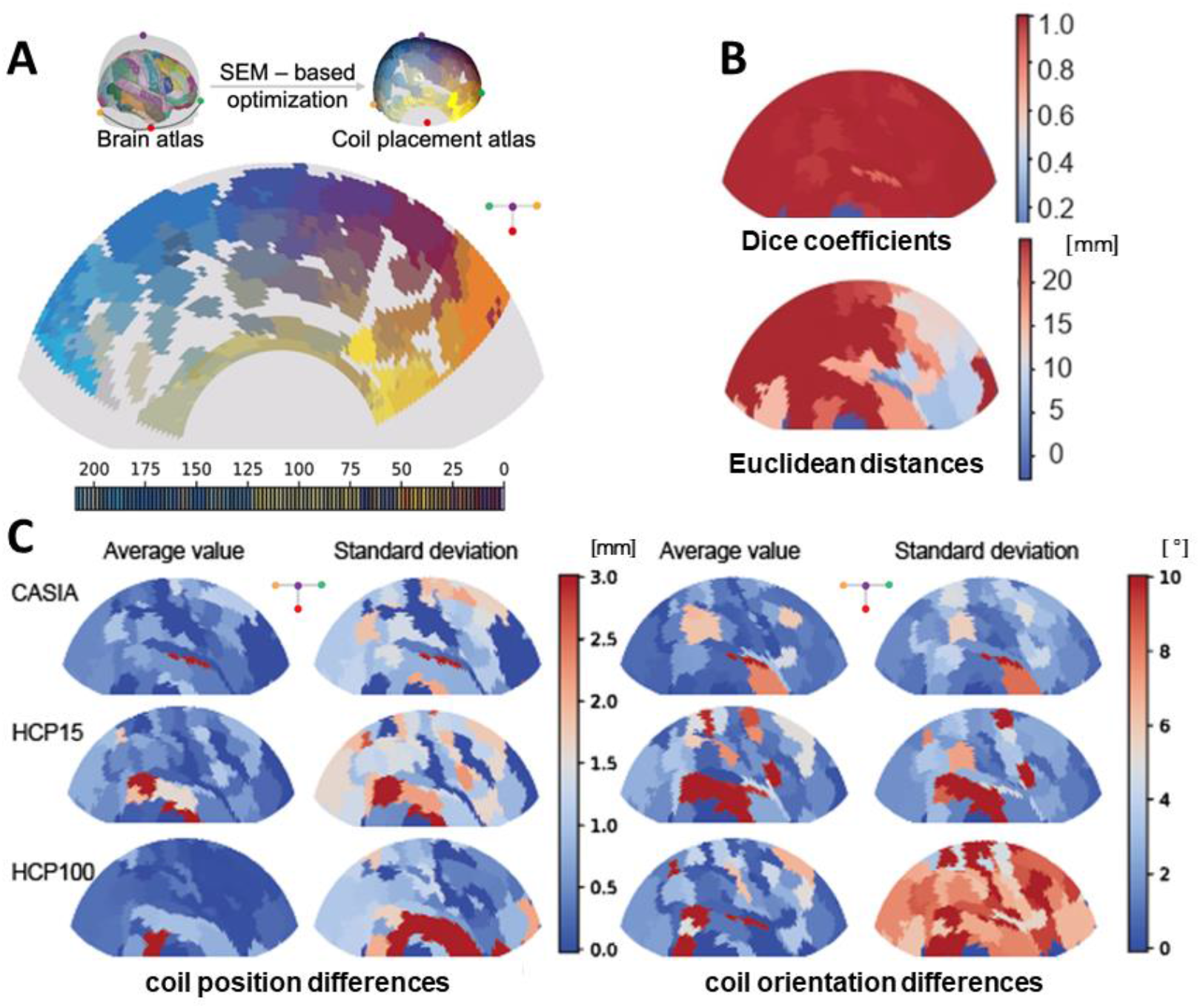
SEM calculation for all 74 target ROIs. Panel A represents the candidate regions for all the target ROIs in the CASIA dataset. Panel B represents the Dice coefficients and the Euclidean distances of the candidate regions for all the 74 target ROIs calculated by the leave-one-subject-out method in the CASIA dataset. Panel C represents the mean values and standard deviations of coil position and orientation differences for all 74 target ROIs in the CASIA, HCP15 and HCP100 datasets.

We compared the Z-scores of E-norm values for the target region using two distinct methods: the SEM framework and the ADM approach. The E-norm values obtained via SEM were more significant than those obtained from the ADM approach in the target region for both the DP and CASIA II datasets (Figure 4A&B).

**Figure 4.**
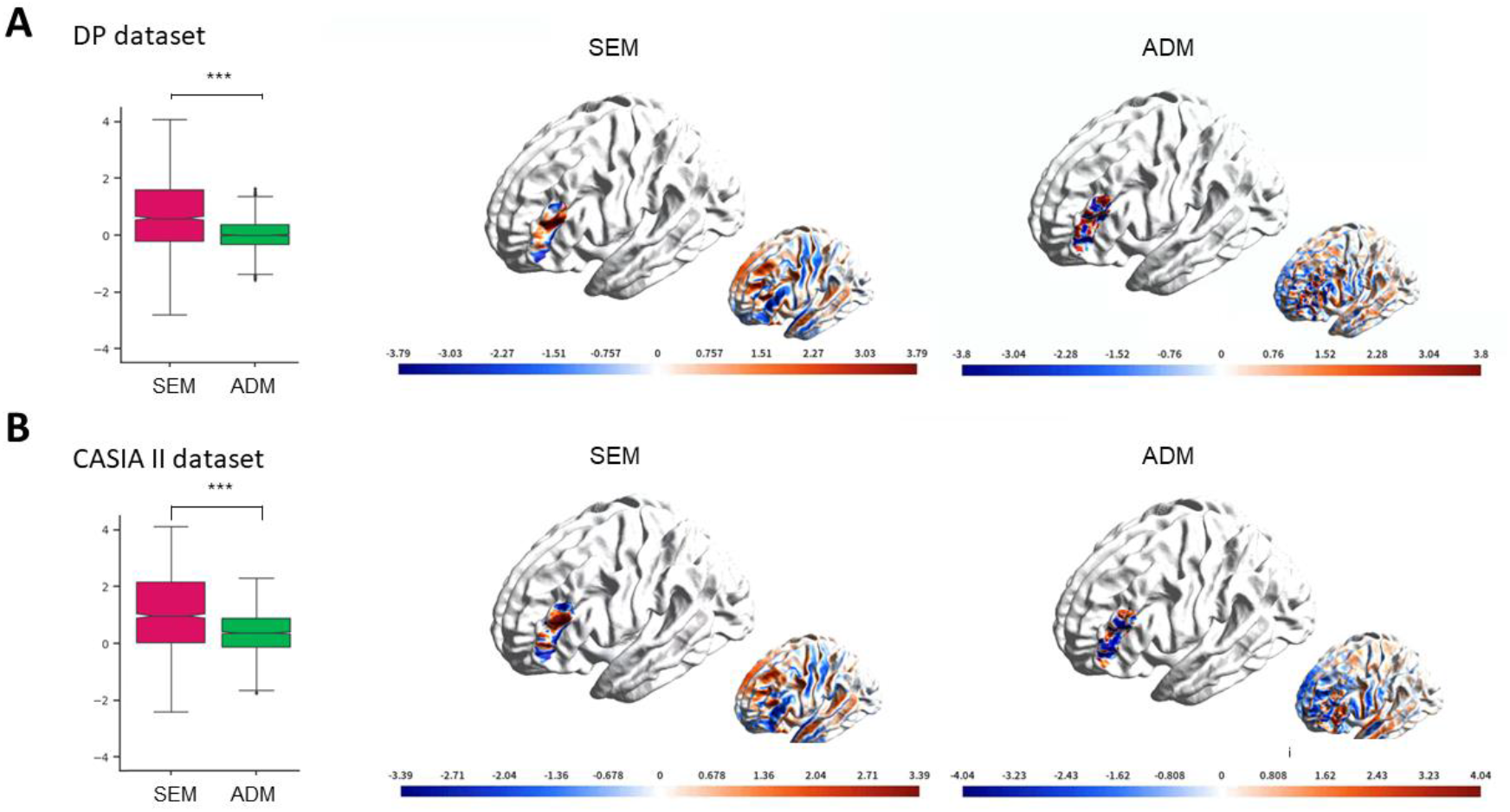
Comparison of E-field normal (E-norm) values for optimal coil placement using SEM and ADM methods. A, Z-scores of the E-norm values obtained using the two methods for the DP dataset. B, Z-scores of the E-norm values obtained using the two methods for the CASIA II dataset.

## Discussion

We suggest using the SEM framework to determine optimal coil placement. The CASIA dataset served as a template to build the SEM framework. We used the SEM framework to give the optimal coil placement in HAC polar and MNI coordinates. The SEM framework was tested for consistency using the CASIA, HCP15, and HCP100 datasets. The E-norm of optimal coil placement discovered by SEM was compared with those found by ADM in the DP and CASIA II datasets. Our findings demonstrated that SEM can achieve optimal coil placement. The E-norm of SEM was more concentrated within the target ROI than other approaches.

The optimal coil location for A9/46v, determined by SEM (HAC Polar coordinates: 41.997°, 120.383°; MNI coordinates: −60 mm, 56 mm, 27 mm), was found to be more lateral and anterior compared to the conventional 5-cm rule. These findings align with previous research by Herbsman et al. (2009), which also indicated a more lateral and anterior coil placement. We have demonstrated consistency for SEM across three independent datasets, employing approximately 5 million simulations. The consistency can be found across different databases and in various target ROIs, which was in line with previous studies that have obtained consistent findings (Balderston et al., 2022; Gomez-Tames et al., 2018; Gomez et al., 2021). Furthermore, it was demonstrated that the SEM framework may improve the effectiveness of coil placement optimization because the E-norm induced by SEM was much larger than that of the ADM technique. The SEM framework also provided prior knowledge of group-level coil placements, as shown in Supplementary Method S2 and Figure S6-8.

We built the HAC polar coordinate system in the SEM framework. Unlike the coordinate system built by Xiao et al. (2018), we used the method of dividing the angles equally. This not only allowed for a more reasonable division of the polar coordinate system. More importantly, the large-scale coil placement based on our coordinate system was more tightly arranged. Cartesian coordinate systems used in conventional scalp-surface-based methods (Jiang et al., 2022; Jurcak et al., 2007; Okamoto et al., 2004; Tsuzuki et al., 2016) often rely on scalp curve length proportions. Those systems are often sensitive to head shapes and perform optimally only when the head is an ideal sphere (Balderston et al., 2022; Balderston et al., 2020; Bungert et al., 2017; Cardenas et al., 2022; Fitzgerald, 2021; Gomez-Tames et al., 2018; Gomez et al., 2021; Herbsman et al., 2009; Jurcak et al., 2007). The HAC system is insensitive to the choice of the initial center point, reducing sensitivity to anatomical variations across individuals. By converting the HAC coordinates of optimal coil placements to MNI coordinates, these placements can be easily integrated into existing commercially available coil localizing devices, such as neuronavigation systems.

The template dataset for this research was MRIs of 18 subjects from the CASIA dataset. Our goal for every subject was to get as many large-scale coil placements (43,116) as possible. When the SEM framework was being built, the E-field simulation of every coil placement was calculated by SimNIBS, which takes a lot of time and processing. In the future, we intend to develop faster E-field calculation methods and more reasonable coil placement selections to enable us to add more subjects’ data as the template. Additionally, we supplied the optimal coil placement for the A9/46v region in the research. This was due to our desire to contrast the coil position with those of Herbsman et al. (2009). In the future, more optimal coil placements for other target ROIs, like M1, can be identified. Not only can prior knowledge of optimal coil placements for motor regions be found, but we can also conduct experiments and compare the coil placements eliciting maximum motor evoked potential with the optimal coil placement calculated by SEM.

## Conclusion

We developed the SEM framework to pinpoint optimal coil placement without time-consuming segmentation or meshing. Our study provided precise coordinates for the optimal coil placement in both HAC and MNI coordinate systems within the target region. Validation across multiple datasets confirmed the consistency of the SEM approach across various target regions. By simplifying the process of determining optimal coil placement, we aim to enhance the convenience and efficacy of TMS assessment and therapy.

## Ethical Approval

All procedures performed in studies involving human participants were by the ethical standards of the institutional and national research committee and with the 1964 Helsinki Declaration and its later amendments or comparable ethical standards.

## Acknowledgments

We are very grateful to all the participants who contributed to this article. R. E. Perozzi and E. F. Perozzi assisted with English language and editing.

## Data and Code Availability

We have made the simulation dataset, constructed using neuroimages from the CASIA I and CASIA II datasets, publicly available as a resource. This dataset is ready for E-fields modeling and can be accessed through Science Data Bank upon acceptance. The HCP dataset is also publicly available for reference. For clinical data, interested parties can request access from the corresponding author, T. J., by submitting a reasonable request. The code used for mapping stimulation effects and constructing the optimal coil placement atlas, along with the data supporting the results of this study, has been uploaded to GitHub. You can find the code and data at the following GitHub repository: https://github.com/ZhongGangliang/TMS_OPA.

## Authors’ Contributions

T.J. proposed the concept and designed the protocol. G.Z. performed the concept and analyzed data. F.J., L.M. and Y.Y. performed the experiments, and B.Z. performed the MRI scanning. D.C., J.L., and N.Z. contributed to the data analysis. T.J., Z.Y., and L.F. led the project and supervised the experiments. All authors contributed to the writing of the manuscript. Data access and verification: G.Z., L.M., and Y.Y.

