## Supplementary Material for "Stimulation Effects Mapping for Optimizing Coil Placement for Transcranial Magnetic Stimulation"

This file includes:

Supplementary material 1: Datasets & TableS1

Supplementary material 2: Supplementary Methods S1 & S2

Supplementary material 3: Supplementary Figure S1 to S9

Supplementary material 4: HAC system

### Supplementary material 1: Datasets

#### CASIA dataset

*Subjects.* The CASIA dataset included 18 healthy, right-handed subjects with no history of neurological or psychiatric disorders (9 males, age range, 23–31 years; age, 25.28 ± 2.05 years). Written informed consent was obtained from all participants before the scans. The MRI data acquisition was approved by the Ethical Committee of the Institute of Automation, Chinese Academy of Sciences.

*Magnetic resonance imaging (MRI) data acquisition*. MRI data was acquired using optimized sequences to ensure accurate reconstruction of head models (Nielsen et al., 2018). Individual MR images were acquired with a GE UHP 3.0 T MR scanner: (i) T1-weighted: 342 sagittal slices, matrix size = 340×340, voxel size = 0.7 × 0.7 × 0.7 mm³, flip angle 15°, TR/TE/TI = 4666/4.66/2200 ms (repetition, spin echo, inversion time), (ii) T2-weighted: 324 sagittal slices, matrix size = 360 × 360, voxel size = 0.7 × 0.7 × 0.7 mm³, flip angle 90° TR/TE/TI = 4002/114/1099 ms, (iii) diffusion MRI (dMRI): 96 axial slices, matrix size = 152 × 152, voxel size = 1.5 × 1.5 × 1.5 mm³, TR/TE 10000/52 ms, flip angle 90°, 60 diffusion directions, b-value 1000 s/mm³. An additional b0 image with the reversed encoding direction was acquired and used during the calculation of the anisotropic conductivity sensors for eddy current correction and distortion correction. Technical details are provided in Supplementary Table S1.

#### HCP15 dataset

*Subjects and MRI data acquisitions.* The HCP15 dataset consisted of MRI data from 15 healthy (10 males, age range, 24–33 years, age, 28.93 ± 2.79 years), and unrelated adults obtained from the Human Connectome Project (HCP) database. These subjects were adopted in this study because the possibility of reconstructing smooth and high-resolution skull and skin surfaces had been verified in (Htet et al., 2018). Subject 209733 was abandoned because of an abnormal segmentation of brain tissues. The T1-weighted, T2-weighted, and dMRI images were downloaded in a preprocessed form, following the minimal preprocessing pipeline. The scanning parameters and preprocessing details were specified in previous studies (Glasser et al., 2013; Van Essen et al., 2012).

#### HCP100 dataset

*Subjects and MRI data acquisitions.* The HCP100 dataset included MRI data from 100 healthy (39 males, age range, 22–35 years, age, 29.05 ± 3.38 years), and unrelated adults obtained from the HCP database. The T1-weighted and dMRI images were used for the repeatability validation of the SEMs and optimal TMS coil placements. The details of the data acquisition were the same as for the HCP15 dataset.

### DP dataset (Depression study)

*Participants.* Participants in the depression study were recruited from the Second Affiliated Hospital of Xinxiang Medical University. Diagnosis of depression was made based on the Structured Clinical Interview for DSM-IV-TR Axis I Disorders. The severity of depression and anxiety symptoms was assessed using the Beck Depression Inventory (BDI) and Beck Anxiety Inventory (BAI). The study protocol was approved by the Ethics Committees of the hospital, and written informed consent was obtained from all participants. A total of 54 subjects (Han Chinese ancestry, 25 males, age range = 18-55 years, age = 34.15 ± 12.02 years) who completed the treatment were involved in the analysis.

*MRI data acquisition.* MRI scans were performed using a Siemens Verio 3.0 T MR scanner. The T1-weighted images used in the current study were collected with 192 sagittal slices, matrix size = 256×256, voxel size = 1 × 1 × 1 mm$^3$, flip angle 7°, TR/TE/TI = 2530/2.43/1100 ms.

*TMS strategies for depression treatment.* All subjects received TMS treatment sessions using a Magstim Rapid2 stimulator and a Magstim 70 mm figure-of-eight stimulation coil, navigated by a commercial BrainSight system (The Rogue Research Inc.). Stimulation was delivered to the A9/46v cortex at a frequency of 10 Hz. Depression severity was assessed using the BDI and BAI before and after the treatment course.

### CASIA II dataset

*Participants and MRI data acquisition.* The CASIA II dataset included 25 healthy, right-handed individuals (14 males, age range, 24–30 years, age, 25.12 ± 1.88 years). Written informed consent was obtained from all participants before the examination. The T1-weighted image was used to simulate the E-fields distribution, and the details of the MRI data acquisition were the same as those of the CASIA dataset. The study was approved by the Ethical Committee of the Institute of Automation, Chinese Academy of Sciences.

**Supplementary Table S1.**

Technical details of the MR scanner, software, and head coils, stratified according to dataset.

| Datasets | Number | Age (y) | Sex (M/F) | MRI<br>Scanner | Head coil |
|---|---|---|---|---|---|
| CASIA | Healthy 18 | 25.28 ± 2.05 | 9/9 | GE<br>Signa UHP 3.0T | 48 channels |
| HCP15 | Healthy 15 | 28.93 ± 2.79 | 10/5 | Siemens | 32 channels |

| | | | | | |
|---|---|---|---|---|---|
| | | | | Skyra 3.0 T | |
| HCP100 | Healthy 100 | 29.05 ± 3.38 | 39/61 | Siemens Skyra 3.0 T | 32 channels |
| DP | Patients 54 | 34.15 ± 12.02 | 25/29 | Siemens Verio 3.0T | 8 channels |
| CASIA II | Healthy 25 | 25.12 ± 1.88 | 14/11 | GE Signa UHP 3.0T | 48 channels |

### Supplementary material 2: Supplementary Methods S1 & S2

### Supplementary Method S1. Reconstructing models for simulating stimulation effects

We used the pipeline in Simulation of Non-Invasive Brain Stimulation (SimNIBS) software [9] or an anatomically realistic volume conductor model of a subject-specific head, as shown in Supplementary figure 1. The individual head models were generated from the MRI data using the SimNIBS headreco command described in [1], employing the Statistical Parametric Mapping (SPM12) Toolbox (http://www.fil.ion.ucl.ac.uk/spm) and Computational Anatomy Toolbox 12 (CAT12) toolbox (http://www.neuro.uni-jena.de/cat). The head models were represented by about 4 million tetrahedra with a vertex density of 0.5 nodes per $mm^2$. T1-or T2-weighted images were used to segment the main tissues of an individual head: scalp, skull, grey matter (GM), white matter (WM), cerebrospinal fluid (CSF), and eyes. All tissues were modelled as isotropic conductors, and the assigned electric conductivity values were 0.465 S/m (scalp), 0.01 S/m (skull), 0.275 S/m (GM), 0.126 S/m (WM), 1.79 S/m (CSF) and 0.50 S/m (eyes). The electric conductivity of the WM could also be assumed to be anisotropic and could be calculated by the conductivity tensors reconstructed by dMRI using the volume normalized mapping approach [10]. Because of differences in the characteristics and quality of the MRI data in the datasets, the model reconstruction and electric conductivity definition of each dataset may differ. The details are provided in Supplementary figure 9. In addition, a figure-eight TMS coil model (corresponding to Magstim 70 mm stimulation coil) [11] provided by the SimNIBS software was used. The coil current was fixed to 1 A for all simulations. A distance of 6 mm was maintained between the center of the figure-eight coil, where the two loops meet, and the scalp point.

### Supplementary Method S2. Optimizing individual coil placement based on a priori group-level optimal coil placement

To develop a general application example of the SEM, we built SEMs for each individual and optimized an individual coil placement based on the group-level optimal coil placement Because a superior coil localization needed to be defined for each subject, an individual optimization strategy based on the group-level optimal coil placement was provided. In our group-level optimal coil placement-based optimization strategy, the intersection regions of the group-level optimal coil placement were used to define an *a priori* range of coil positions and orientations. Based on the coil placements provided by the group-level optimal coil placement, we constructed an envelope surface to provide a continuous HAC parameter of the coil locations $(\alpha, \beta)$ for the group-level optimal coil placement-based optimization strategy. To test the performance of the coil placements priors in the optimization, we compared them with a searching range commonly used in traditional optimization strategies. The searching range imitates the range of the square grid used for motor mapping [7].

Similarly, we constructed an envelope surface based on a square range over the scalp surface with a length of 25 mm to provide a continuous HAC parameter of the coil locations for the traditional optimization strategy. In addition, the searching ranges of the coil orientation in these two optimization strategies were the same, and the optimization range of the coil orientation was from 0 to 180 degrees. Then, both optimization strategies used the same Nelder-Mead (NM) algorithm to optimize the coil placement ($\alpha,\ \beta,\ \theta$). We used the NM algorithm because it is a derivative-free optimization method, has a simple calculation scheme, and enables a strong local search, so it has been widely used [8]. Optimizing the coil placement of a targeted ROI can be transferred to reveal the maximum of the objective function (eq. 1). The objective function is defined as the difference in IFV between the first coil placement and the last coil placement of the targeted ROI, which has three variables, $\alpha$, $\beta$, and $\theta$.

$$Max\, F, F \coloneqq infocEF_{(\alpha,\beta,\theta)\in m}(\alpha,\beta,\theta) \tag{1}$$

where $m$ is related to the prior range of coil placements. The initial coil placement of our optimization strategy is the optimal coil placement of the corresponding target brain, and the center of the range of the coil positions and a 0º orientation were used in traditional individual optimization strategies.

### Supplementary material 3: Supplementary Figure S1 to S9

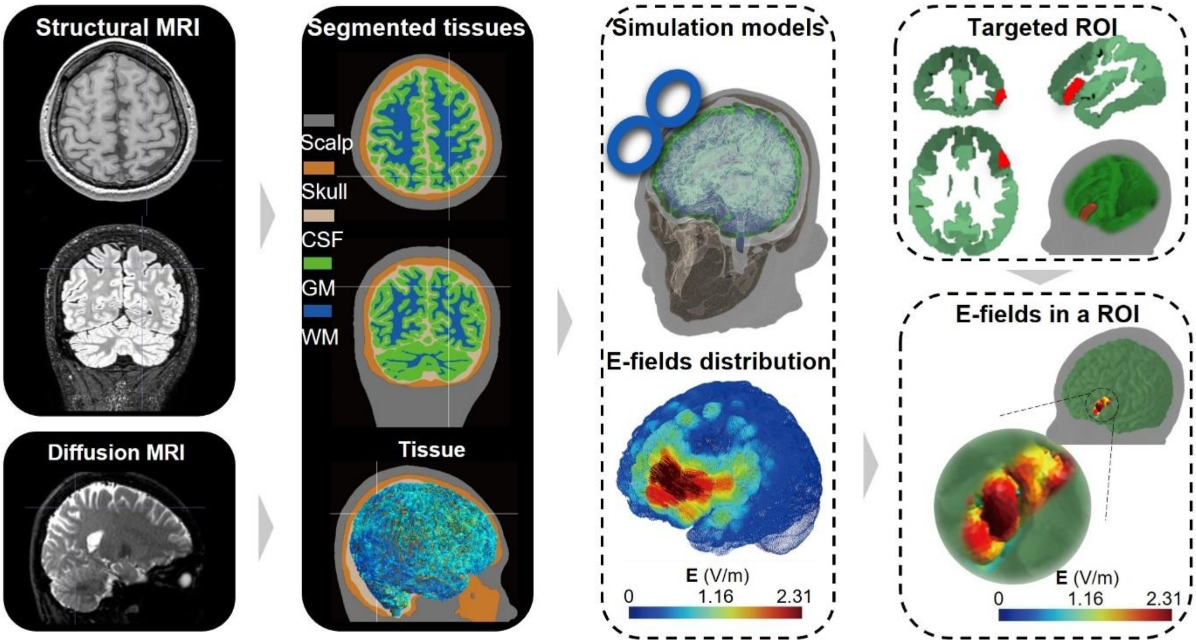


**Figure S1.** Framework for building the simulation model and the distribution of simulated E-fields on the brain and a targeted region of interest (ROI). The simulation model was created using the finite element method (FEM), which involved constructing a realistic head model from magnetic resonance images (MRI) of a healthy subject. The FEM model was then used to simulate the E-field distribution generated by the TMS coil at different placements. The simulated E-field was subsequently normalized using the z-score method, and the distribution of E-fields in the targeted ROI was calculated.

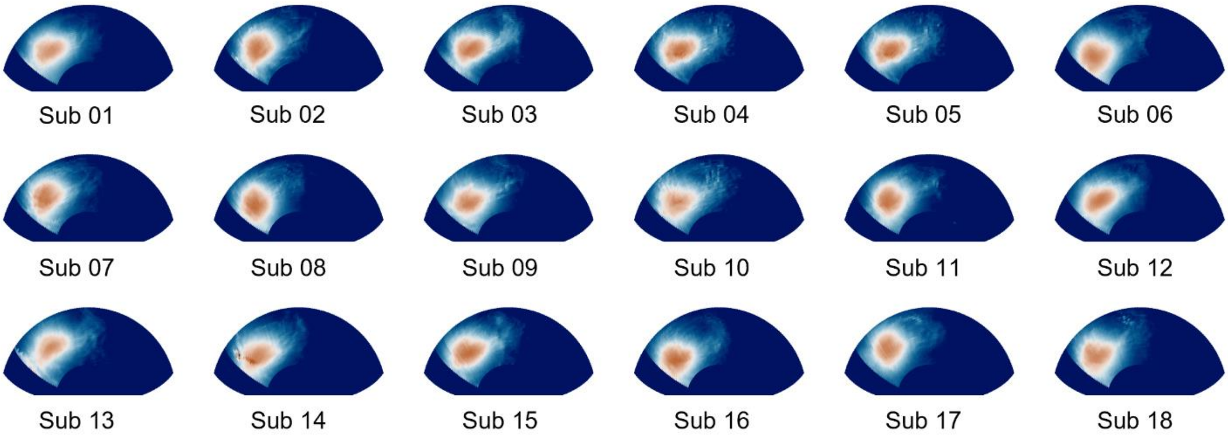


**Figure S2.** Distributions of IFV of the targeted A9/46v subregion for all subjects (*n* = 18) in the CASIA dataset.

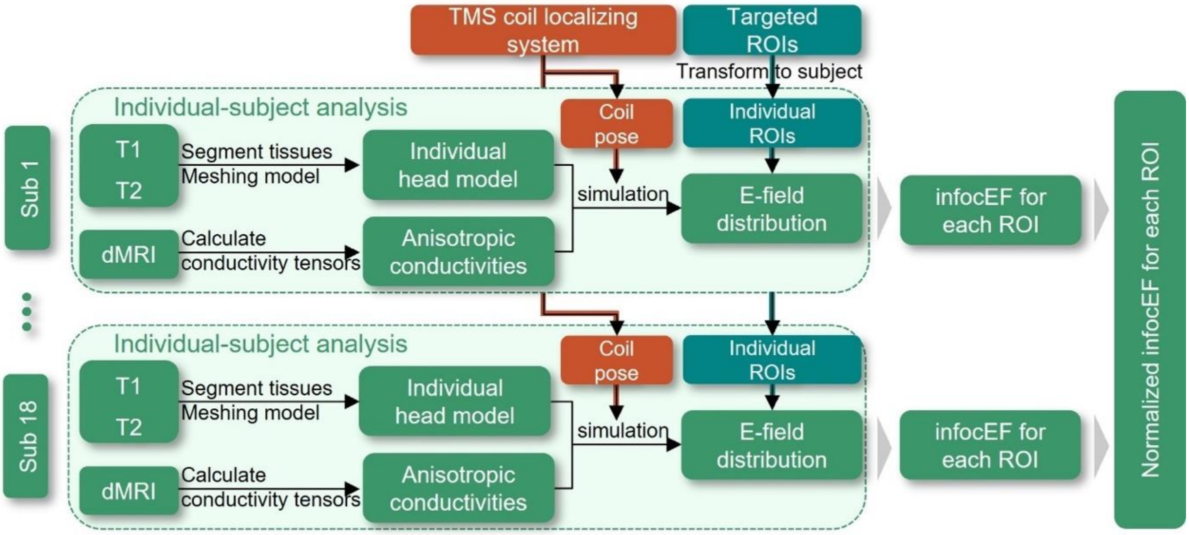


**Figure S3.** Group-level analysis of the IFV of the stimulation effects. For each individual, we constructed an individual head model with anisotropic conductivities based on T1, T2, and dMRI data. Using the TMS coil localizing system, we simulated the E-field distribution based on the coil placement. We then obtained IFV for each targeted ROI. Finally, we normalized the IFV for all subjects using the z-score method to perform group-level analysis.

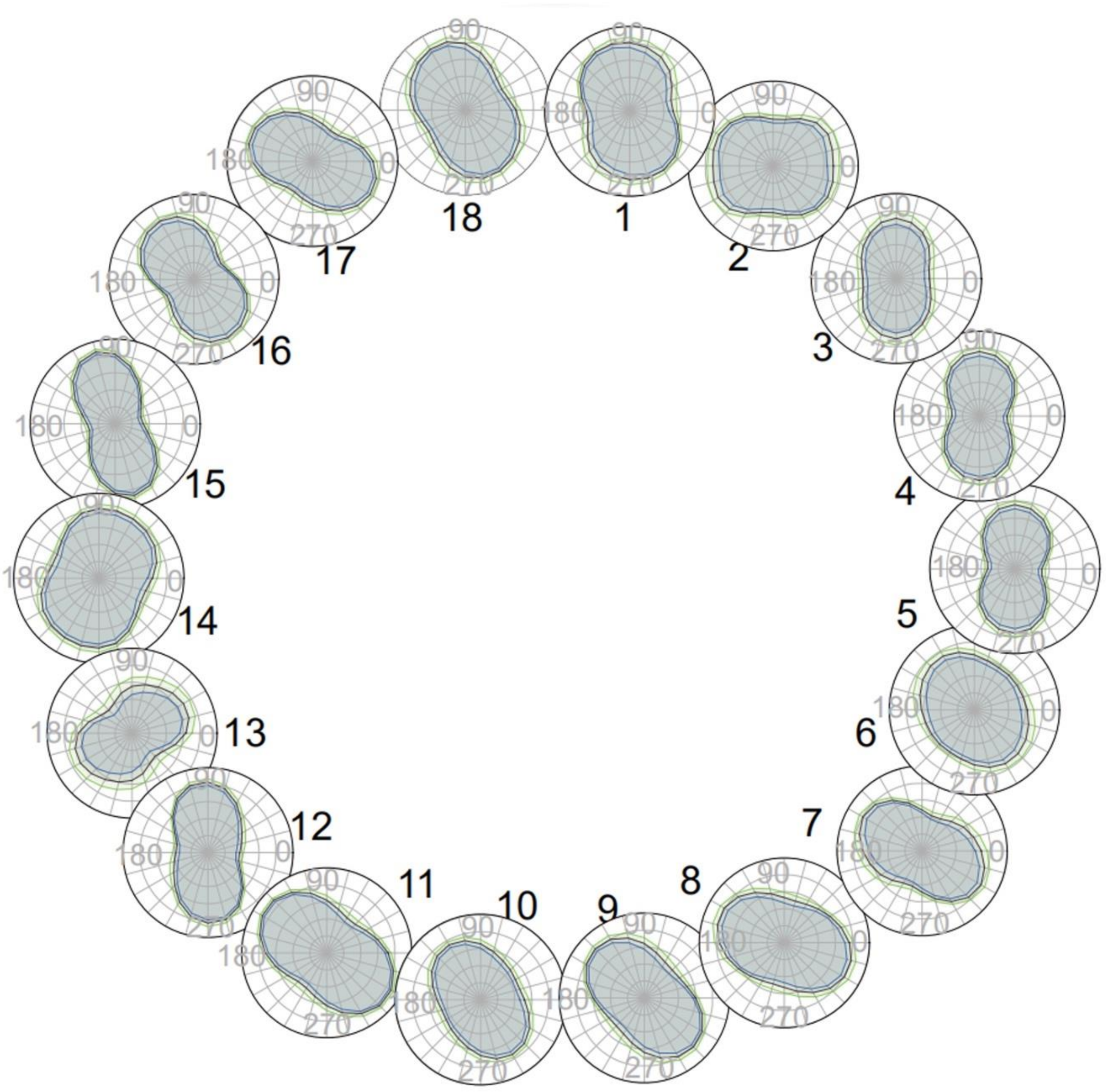


**Figure S4.** Candidate coil orientations of the targeted A9/46v subregion for all subjects (n = 18) in the CASIA dataset.

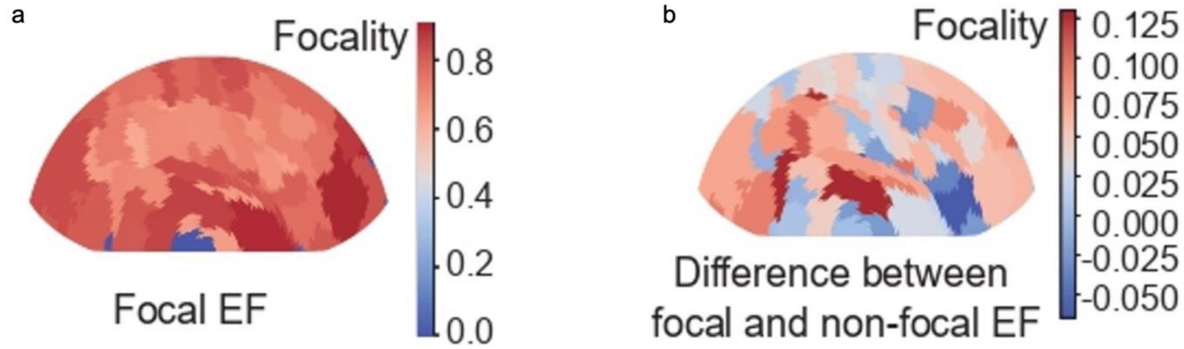


**Figure S5.** a, the focality of the aggregative coil placements was evaluated for 74 targeted ROIs (For a detailed definition of the focality score, please refer to Materials and Methods, "Focality score for population consistency" in Statistics.). b, the difference between the focality scores for the aggregative coil placements.

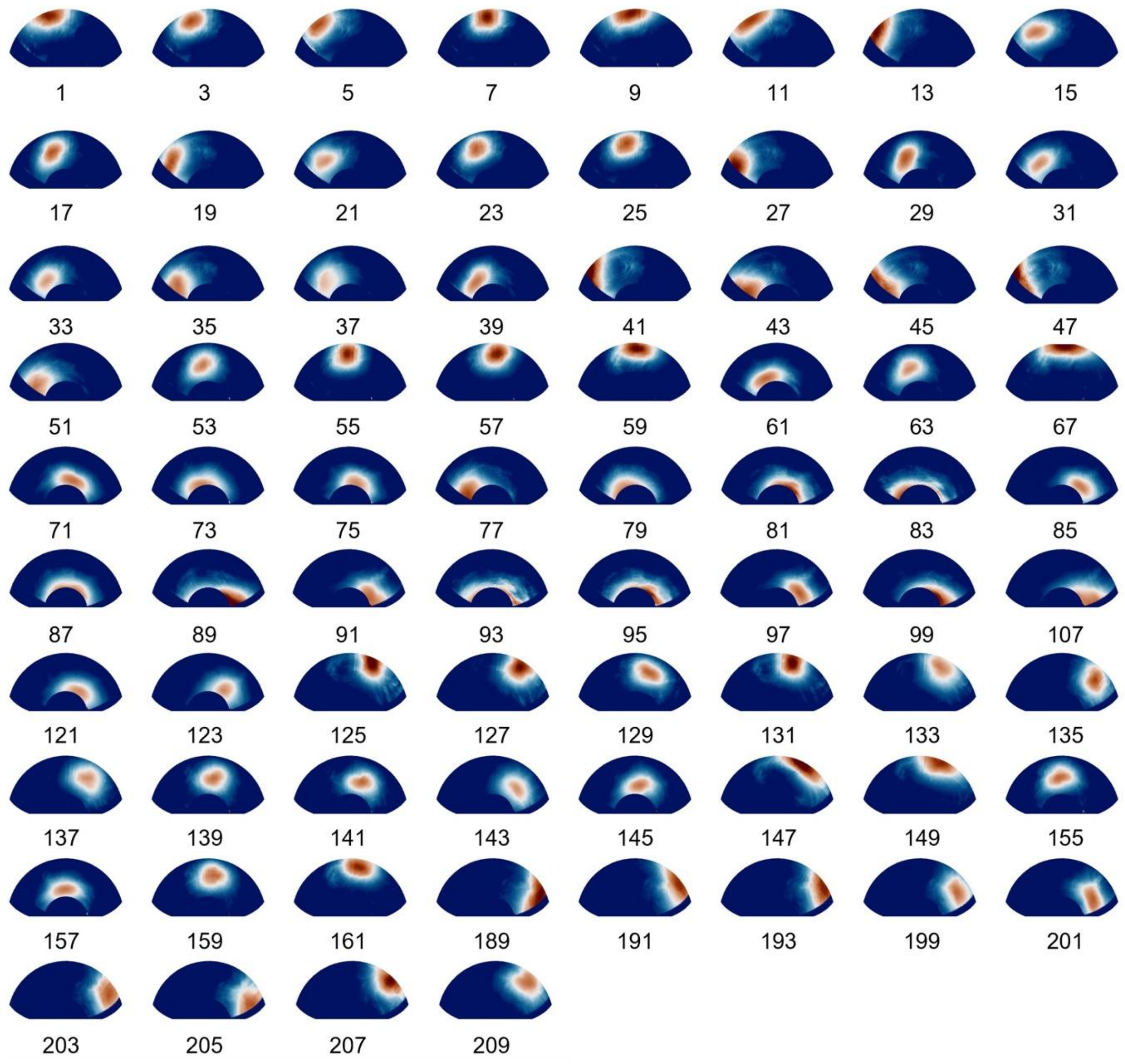


**Figure S6.** SEMs of 74 targeted ROIs for the first subject in the CASIA dataset.

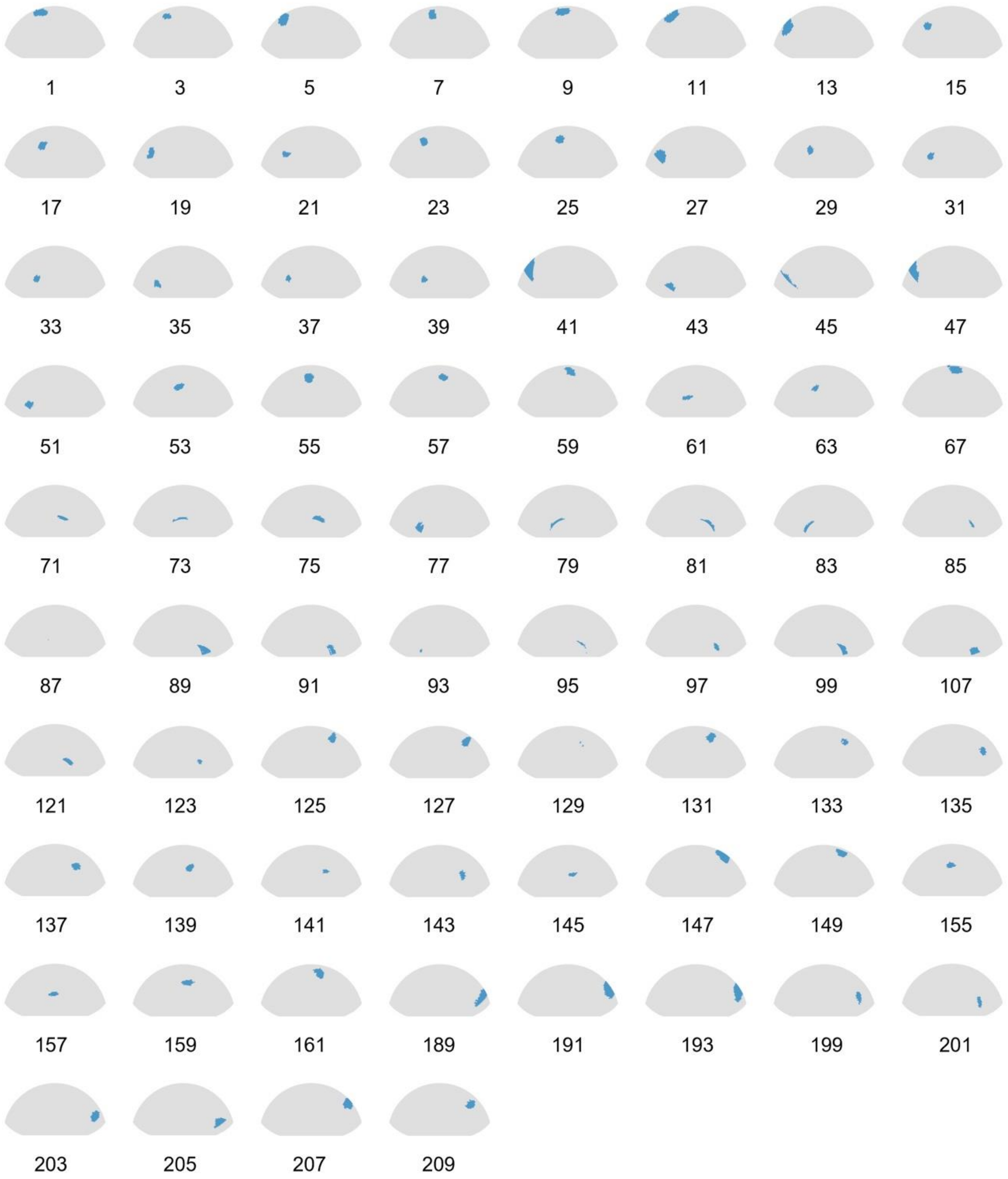


**Figure S7.** *A priori* information about optimal group-level coil positions.

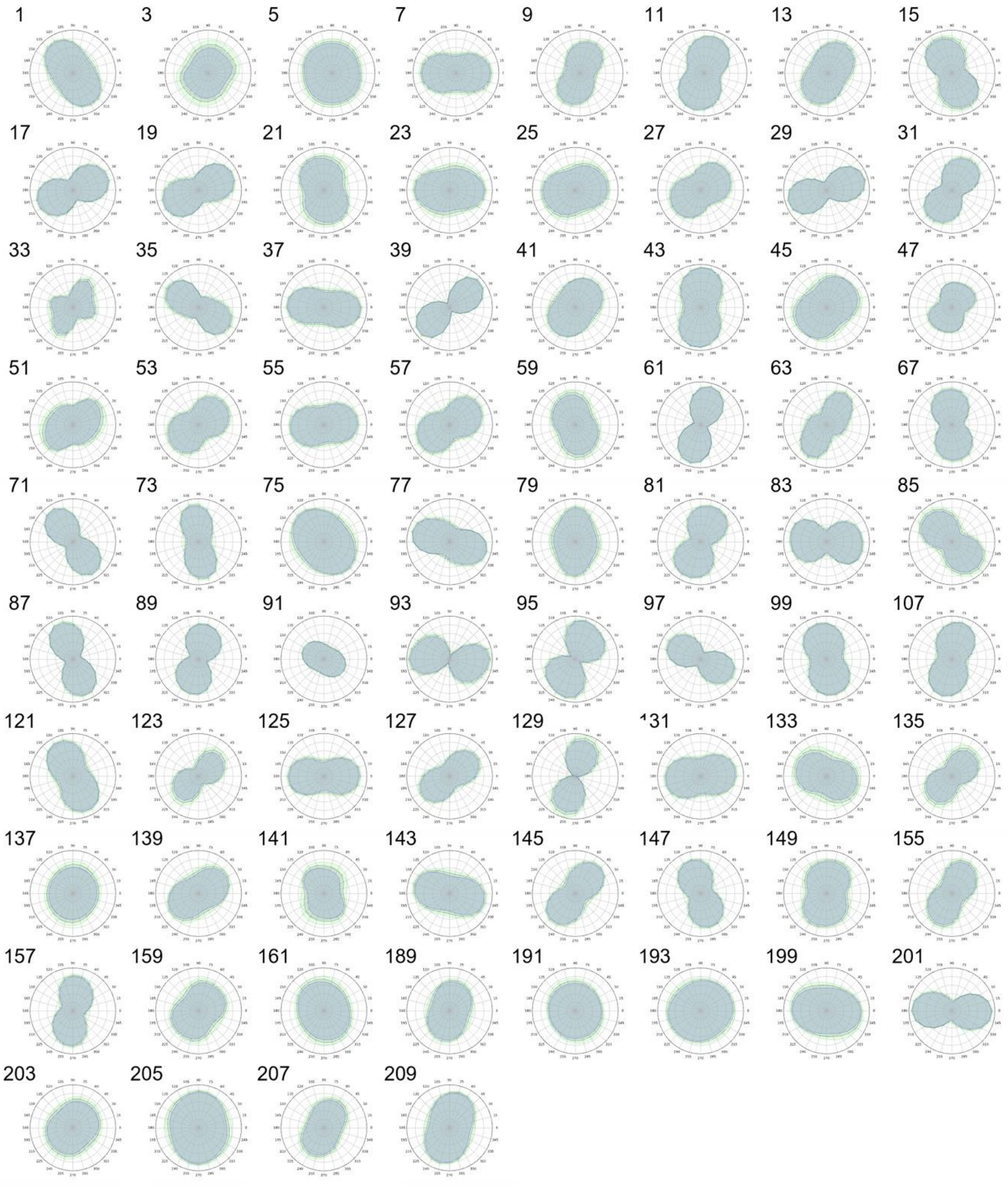


**Figure S8.** *A priori* information about optimal group-level coil orientations .

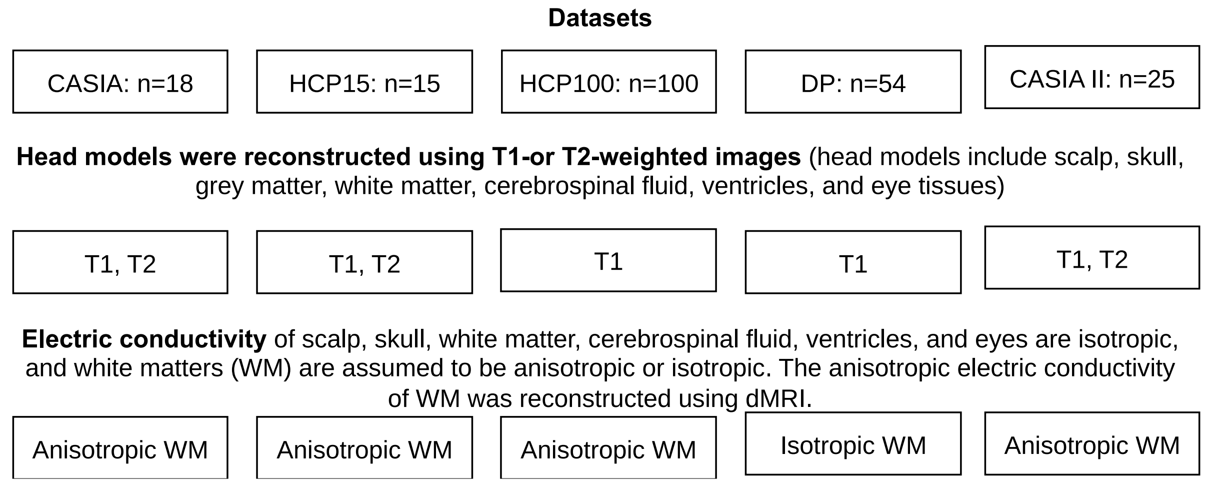


**Figure S9.** Details of model reconstruction and electric conductivity definition for each MRI dataset used in the study.

### Supplementary material 4: HAC system

The HAC system (see Figure S10-S11) was designed to describe the large-scale coil placements on the head. First, five cranial landmarks were inherited from the international 10-20 system (Klem et al., 1999): the nasion (Nz), inion (Iz), left/right preauricular point (LPA/RPA), and Cz. The positions of the landmarks were calculated based on a population average head model reconstructed by 152MNI template images. Then, these landmarks were transformed into individual MRI and assigned to the nearest point on the scalp. Next, the spherical polar coordinates were defined in the HAC system. In the spherical polar coordinate system, O was represented by the origin point in space; the *x*-axis was represented by the orientation of O to LPA, the orientation of *O-Iz* represents the *y-axis*, and the *z*-axis was represented by the O-Cz orientation with the origin at the midpoint of the RPA-LPA line. $\alpha$ was the polar angle in the *x-y* plane from the *z*-axis with $0 < \alpha < \pi$, and $\beta$ was the azimuthal angle in the *x-y* plane from the x-axis with $0 < \beta < \pi$ of the HAC system. Finally, the positions on the scalp surface were parametrized as continuous angle coordinates ($\alpha$, $\beta$) based on the tetrahedrons of the head model.

Based on the HAC system, unlike neuroimage-based strategies that determine coil placement using position (x, y, z coordinates) and orientation (roll, pitch, and yaw) (Balderston et al., 2020; Gomez-Tames et al., 2018), we parametrize the coil placements using angle coordinates (α, β, θ) representing the coil position (α, β coordinates) and orientation (θ coordinate) in the first step of SEM (see Figure 3 for more details). This reduced the six to three degrees of freedom of angle coordinates, simplifying the quantitative description of coil placements. The TMS coil orientation was defined as the coil handle orientation when the coil was tangential to the scalp site, ranging from -90 to 90 degrees. In our case, (i) the angle between the O-Nz line and O-Iz line was separated into 100 equal parts to determine points $p_i$ ($i$=1, …, 101) over the scalp, then the number of equal parts was scaled to adjust for the real need. (ii) For the left scalp, the angle between the O-LPA line and the O-$p_i$ line was separated into 50 parts to determine points $p_{ij}$ ($j$=1, …, 51), and the right scalp was done similarly. The coil orientation ($\vartheta$) was defined as the coil handle orientation when the coil was perpendicular to the location on the scalp. Overall, coil placement can be parametrized as ($\alpha$, $\beta$, $\vartheta$) based on the HAC system.

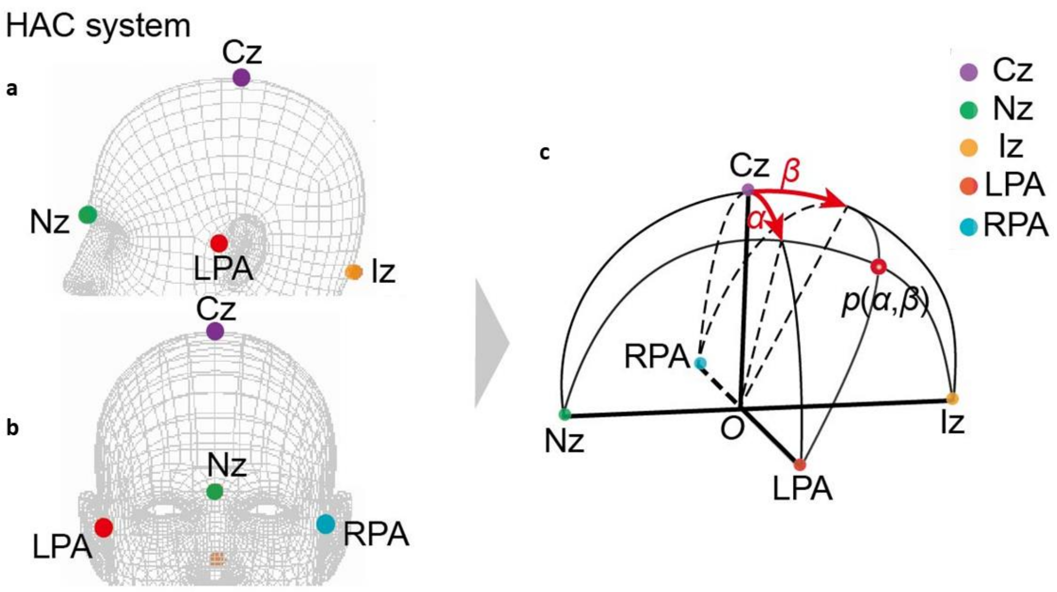


**Figure S10**. Schema of the head-anatomy-based coordinates (HAC) system. The landmarks in a and b show the five cranial landmarks (Nz, Iz, LPA, RPA, Cz) constructed in the HAC coordinate system similar to the 10-20 system (Jiang et al., 2022; Jurcak et al., 2007; Okamoto et al., 2004; Tsuzuki et al., 2016; Xiao et al., 2018). The angle coordinates (α, β) in c represent the TMS coil position on the scalp.

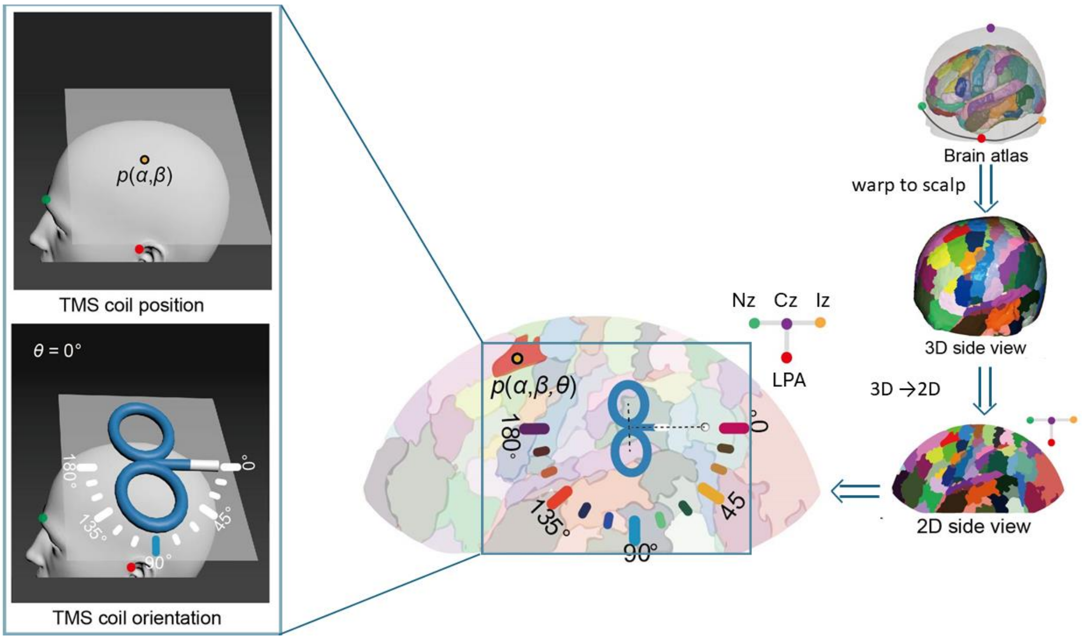


**Figure S11**. The central panel displays the coil placement according to the head anatomy-based coordinates (HAC) system coordinates. The placement of the coil was determined based on the position and orientation of the TMS coil, as shown in the left panel. The 2D representation of the middle cortex was derived from the inflated 3D cortex obtained from a brain atlas in the right panel.

1. Nielsen JD, Madsen KH, Puonti O, et al. Automatic skull segmentation from MR images

for realistic volume conductor models of the head: Assessment of the state-of-the-art. *Neuroimage* 2018; **174**: 587-98.
